# Ambiguity coding allows accurate inference of evolutionary parameters from alignments in an aggregated state-space

**DOI:** 10.1101/802603

**Authors:** Claudia C. Weber, Umberto Perron, Dearbhaile Casey, Ziheng Yang, Nick Goldman

**Author notes:** Smurfit Institute of Genetics, Trinity College, Dublin 2, Ireland (Dearbhaile Casey).

## Abstract

How can we best learn the history of a protein’s evolution? Ideally, a model of sequence evolution should capture both the process that generates genetic variation and the functional constraints determining which changes are fixed. However, in practical terms the most suitable approach may simply be the one that combines the convenience of easily available input data with the ability to return useful parameter estimates. For example, we might be interested in a measure of the strength of selection (typically obtained using a codon model) or an ancestral structure (obtained using structural modelling based on inferred amino acid sequence and side chain configuration).

But what if data in the relevant state-space are not readily available? We show that it is possible to obtain accurate estimates of the outputs of interest using an established method for handling missing data. Encoding observed characters in an alignment as ambiguous representations of characters in a larger state-space allows the application of models with the desired features to data that lack the resolution that is normally required. This strategy is viable because the evolutionary path taken through the observed space contains information about states that were likely visited in the “unseen” state-space. To illustrate this, we consider two examples with amino acid sequences as input.

We show that *ω*, a parameter describing the relative strength of selection on non-synonymous and synonymous changes, can be estimated in an unbiased manner using an adapted version of a standard 61-state codon model. Using simulated and empirical data, we find that ancestral amino acid side chain configuration can be inferred by applying a 55-state empirical model to 20-state amino acid data. Where feasible, combining inputs from both ambiguity-coded and fully resolved data improves accuracy. Adding structural information to as few as 12.5% of the sequences in an amino acid alignment results in remarkable ancestral reconstruction performance compared to a benchmark that considers the full rotamer state information. These examples show that our methods permit the recovery of evolutionary information from sequences where it has previously been inaccessible.

## Introduction

The evolution of protein sequences is driven by a combination of forces that influence both what types of mutation occur and which of them are allowed to fix by natural selection. The former process operates at the level of the nucleotide sequence, and manifests at the amino acid level through the structure of the genetic code. Functional and structural constraints then determine the probability of survival of the mutants. A wide variety of computational tools to make inferences about different layers of this process are available, considering observations from nucleotide, codon or amino acid sequences, and (occasionally) protein structure. Models that take data from one of these state-spaces as input typically use transition probabilities between these same character states to compute outputs, such as phylogenies, selective constraints, or ancestral states. Being able to obtain certain types of information about evolution is therefore usually contingent on having access to observations in the relevant state-space.

Given the abundance of available genome sequences, access to interesting data is ordinarily not a problem. Codon sequences, for example, are commonly used to quantify the strength of natural selection, measured by *ω*, the relative rate of non-synonymous to synonymous substitutions. Variants of the standard codon model estimate constraints on specific sites, branches, or different types of amino acid substitutions [1–3]. Empirical amino acid models, which work with amino acid sequences and are often used to estimate phylogenies, consider how “exchangeable” different residues are [4–6]. This allows them to capture some functional constraints and therefore reconstruct plausible amino acid trajectories and ancestral sequences.

A subset of models go beyond sequence alone and incorporate elements of structure. Recently, an extended version of the empirical amino acid model was introduced that additionally accounts for rates of exchange between amino acid side chain configurations [7]. How suited a given amino acid is to a particular sequence and structural context is not only influenced by the biochemical properties of its side chain, but also by its spatial orientation. This includes the rotation of the *C*^*α*^-*C*^*β*^ bond, or the *χ*_1_ rotational isomer (‘rotamer’) configuration, which can be discretized into up to three states per residue, resulting in a state-space with 55 characters (see [7] for details). The empirical rotamer-aware model therefore allows reconstruction of ancestral amino acid sequences that include side chain configurations [7] — provided structural information is available for the extant descendants of the protein of interest. Reconstructed side chain configurations are of practical interest as they can provide a plausible prior for structure prediction for a variety of applications. For example, structural models are widely used for *in silico* functional annotation of genes and variants, prediction of protein-protein interactions and docking [8–10].

Given the variety of options, the choice of model for a study might be guided by the research question and which aspects of the evolutionary process are most interesting. In some cases, data availability may limit the range of suitable models. For example, given a set of amino acid sequences, models operating in codon- or rotamer space cannot be straight-forwardly applied. Attempts to bridge the gap between available input and desired model output have, thus far, been limited — at least as far as conventional observable character states are concerned. For example, Yang et al. [1] formulated an amino acid model that merges synonymous codons into a single amino acid state, with substitution rates computed as an average of the codon rates. This model can estimate transition-transversion bias from amino acids. However, it is unable to provide a measure of the strength of selection (ref. [1], p. 1608).

Nevertheless, there are well-established methods that allow the handling of data for which only part of the state-space information is available. This is achieved by encoding observed states as ambiguous representations of characters in a larger state-space. This application of standard statistical theory for missing data has been used previously in phylogenetics (e.g. ref. [2], p. 110–112), a notable example being covariotide models, where each nucleotide may be in an “on” or “off” state that cannot be directly observed [11]. Can the principles employed by these methods be applied more broadly to allow substitution models to take “partial” observations as input?

In this paper, we demonstrate that it is possible to infer information about evolutionary processes that occurred in an expanded state-space using only the aggregated data, taking advantage of an established method for handling ambiguity in sequence alignments. The ability to model sequences in a state-space with a larger set of characters allows us to obtain outputs that would otherwise be unavailable. For example, we can capture relative selective constraints on non-synonymous versus synonymous substitutions from amino acid sequences. The path through amino acid space hence helps reveal the path evolution takes through codon space. The same method can be applied to reconstructing ancestral amino acid rotamer configurations using only amino acid sequences. Using input data consisting of a mixture of rotamer and amino acid sequences further allows us to refine these reconstructions and obtain a useful starting point for homology modelling.

## Methods

### Inferring model parameters from data in an aggregated state-space using ambiguity

Our framework is maximum likelihood (ML) inference on phylogenetic trees, based on alignments of observable characters that evolve independently according to a Markov process. We consider cases where the characters at the tips of a phylogenetic tree are only available in an ‘aggregated’ state-space **A** with *m* states. Each state *a*_*i*_ in **A** = {*a*_1_, …, *a*_*m*_} corresponds to one or more ‘separate’ states *s*_*j*_ in a larger state-space **S** = {*s*_1_, …, *s*_*n*_} (where *n* > *m*). Meanwhile, each state in **S** maps to a single state in **A**. For example, where **S** describes the set of 61 sense codons, **A** might describe the 20 amino acid states: each codon codes for one specific amino acid, while a given amino acid can be represented by multiple codons. Similarly, each amino acid (**A**) can represent multiple rotamer configurations (**S**; see [7], Table 1). If we only have access to amino acid sequences rather than codon or rotamer sequences, but modelling the data in **S** would be more informative, we can take advantage of these mappings.

In order to estimate phylogenetic models under ML when the data do not match the model state-space, we modify an established method for handling alignment gaps and ambiguity characters. The conditional probability vector *L*_*k*_(*j*) is a crucial part of phylogenetic likelihood calculations. It records the probability of the observed data descended from node *k* conditional on the presence of state *j* at node *k*. There is one such vector for each combination of alignment position (not indicated in this notation, for simplicity) and node *k*, with one element for every permitted state *j*. The iterative calculation of the likelihood is initialised at the tips of the tree: if *k* is a tip with state *x* recorded in the alignment, the element *L*_*k*_(*x*) is set to 1 and *L*_*k*_(*j*) = 0 for all other *j* ≠ *x* [12] — the data are recorded as having been correctly observed with certainty. When data are missing, or when there is a gap at a site in the alignment, the corresponding *L*_*k*_(*j*) are set to 1 for all *j*, representing total absence of knowledge of the true character states and effectively removing node *k* from the likelihood calculation for the site.

In the case where the observed data are in the aggregated state-space **A**, and we are interested in modelling in the separate state-space **S**, we can proceed in a similar manner. Consider a simple four-state (**S**) model with the aggregated (**A**) states *a* = {*a*_1_, *a*_2_} and *b* = {*b*_1_, *b*_2_}. If we observe state *a* in **A**, which could represent either *a*_1_ or *a*_2_ in **S**, the corresponding conditional probability vector 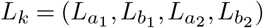 is set to (1, 0, 1, 0). Hence, our observation is ambiguous with respect to the character in **S**. We use the term ‘ambiguous’ to refer to instances where incomplete information about the state at a given site is available, but the character is not missing. Where data are completely absent (missing) for an alignment position, the same vector is encoded by *L*_*k*_(*x*_*j*_) = (1, 1, 1, 1). Once *L*_*k*_ has been set at all tips according to this modification, the calculation of the likelihood proceeds as normal following Felsenstein’s pruning algorithm 13].

Treating data observed in **A** as ambiguous states in **S** is similar to the “covariotide” model of Huelsenbeck [11], which assigns each nucleotide an ambiguous “on” or “off” state. Ambiguity has also been used to encode population allele frequencies using small samples as input [14], and to handle sequence error and uncertainty [15]. Our approach differs from that presented in [1], where all synonymous codons for an amino acid are combined into one state and substitution rates between amino acids represent averages over codons.

### Description of the codon model

The codon model considered here follows the standard M0 model as implemented in PAML ([16]; see also [17,18]), with parameters *ω* = *dN/dS*, *κ* representing the ratio of transition mutations to transversions, and *π* representing the codon equilibrium frequencies. The instantaneous rate matrix is given by:

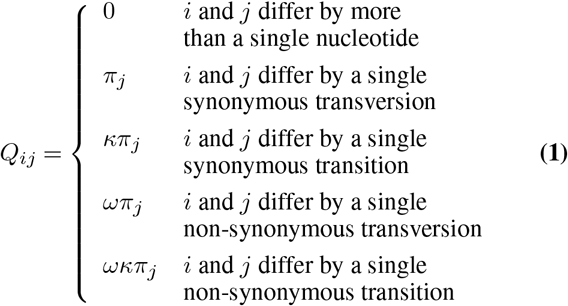

We implemented the model (now available in PAML as M5) and likelihood calculations in a standard ML framework, encoding the conditional probability vector *L*_*k*_ in codon space (**S**) using observed amino acids (**A**), as outlined above. We make the simplifying assumption that the vector of equilibrium frequencies is fixed at *π*_*j*_ = 1*/*61 for all codons *j*. The codon frequencies cannot be directly observed, and examining their identifiability is beyond the scope of this work.

### Codon sequence simulations and inference

We used evolver [16] to generate a single random unrooted tree with 20 tip nodes using a birth-death process [19] with a tree height of 0.5 (see Figure S1A). We next simulated sequences under M0 over a range of parameters generating 100 replicates with 3000 codons for each combination of configurations (unless stated otherwise). We then analysed the simulated sequences using codeml from the PAML package, fitting M5 to the translated amino acid sequences and fitting M0 to the original codon sequences, both assuming equal codon frequencies [16]. We also recorded the standard errors for the parameters (option getSE = 1).

### Description of the rotamer model

The empirical rotameraware model (RAM55) follows the structure of a standard empirical amino acid model and is fit to alignment data in the same manner. The instantaneous rate matrix, *Q*_*ij*_, is defined by the exchangeabilities between states derived from a database of sequences of proteins with known 3D structure and their equilibrium frequencies. Rather than the usual 20 amino acid states, the rotamer model considers 55 discrete states determined by the *χ*_1_ dihedral angle between the side chain’s first two covalently linked carbons, the rotamer configuration. Each amino acid can be categorised into up to three distinct states based on an observed protein structure; for example, state L3 denotes a leucine residue in conformational state 3. RAM55 is described in detail by [7], along with RUM20, a conventional 20-state empirical amino acid model computed from the same dataset.

### Rotamer sequence simulation and ancestral side chain configuration reconstruction

To generate sequences in rotamer space with known phylogenies and ancestral states, we used a set-up similar to the one described by [7]. Briefly, we randomly generated a 32-tip tree using a Yule process and scaled the branches by 0.1–1 (Supplementary Figure S2). We then performed a continuous-time Markov chain simulation along the branches for 1000 replicates each of 200 sites using the RAM55 exchangeabilities and equilibrium frequencies. Simulated alignments use a custom encoding format which expresses both amino acid states and rotamer states (that is, a mixed alignment) using a common alphabet of single-character symbols (see https://bitbucket.org/uperron/ambiguity_coding).

To emulate cases where structural information in not available for some of the terminal nodes, we generated mixed alignments by ‘masking’ a proportion of the terminal rotamer sequences, leaving only amino acids. Amino acid states are then treated as ambiguous rotamer state assignments in the inference step (see below). Further, sequences can be removed from the simulated alignment to illustrate the loss of information caused by discarding sequences entirely when no structure is available. This is done by replacing a specific proportion of the alignment’s sequences with gap characters. Both the masking and discarding operations are performed over a set of sequences selected independently for each replicate according to a uniform distribution.

To reconstruct ancestral states we modified the approach described by Perron et al. [7], encoding each amino acid (statespace **A**) observed in the alignment as ambiguous in rotamer space (state-space **S**) in the conditional probability vector. This procedure allows us to use the RAM55 model to infer rotamer sequences at internal nodes using amino acid or rotamer sequences and the tree that was used for simulation as input. To compute posterior probabilities for reconstructions [20], we applied the marginal reconstruction algorithm of Koshi and Goldstein [21]. A joint reconstruction algorithm [22] gives qualitatively similar results. To assess the accuracy of the reconstructions, we examined the proportion of sites with matching characters in rotamer space. That is, we require both the amino acid and its rotamer configuration to be identical.

The same reconstruction approach can be modified to predict side chain configurations for extant homologous proteins in a given family (i.e. tip nodes of the tree). Specifically, each terminal rotamer sequence is, in turn, masked and its side chain configurations are reconstructed conditioned on the observed amino acid states by treating the terminal node as if it were an internal node. This permits us to infer rotamer configurations for extant proteins with known sequences but unknown structures, based on the known sequences and structures of their homologs. To illustrate this we considered two manually curated empirical datasets, consisting of 16 ADK structures and 30 RuBisCO structures from PDBe [23], respectively. For each dataset, a multiple amino acid sequence alignment was generated using MAFFT [24]. Rotamer configuration was then assigned to each amino acid in the alignment, generating a rotamer sequence alignment (see [7] Methods for details). The tree for the reconstruction was estimated from the rotamer sequence alignment using RAxML-NG under the RAM55 model [7, 25]. We then masked, in turn, each terminal rotamer sequence in the alignment and predict each amino acid’s *χ*_1_ configuration using RAM55 and the marginal reconstruction algorithm as described above. Here, the extant amino acid sequence is known and the rotamer state prediction is thus constrained to the observed amino acid. Prediction accuracy can be computed against the original rotamer sequence; in order to benchmark our method’s accuracy we first established a baseline accuracy by assigning the *χ*_1_ configuration either according to a uniform probability distribution (we denote this by ‘Unif’) or using the relative equilibrium frequencies of each possible configuration according to RAM55 (‘RelFreq’).

A widely used strategy to predict side chain configurations in unresolved structures consists of assigning to each amino acid the same configuration found at the corresponding site in the nearest homologous neighbour’s structure [10, 26]; we refer to this approach as ‘Nearest Neighbour Configuration’ (NNC). NNC is only applicable to sites where the amino acid is conserved in the template, and so our implementation of NNC falls back to a RelFreq strategy for non-conserved sites. We also evaluated a scenario where no structural information is available for the nearest sequence (‘Masked Nearest Neighbour’, MNN). Here, RAM55 can make use of mixed data (both amino acid-only and rotamer sequences) using the ambiguity coding described above.

## Results

### Substitutions in the aggregated state-space contain information about the path taken through the larger state-space

We first consider how it might be feasible to extract information about a process operating in a separate state-space **S** from data in **A**. This will naturally depend on the relationships between the structures of the state-spaces. Where some states in the aggregated space are only accessible via multiple steps in the separate space, it is possible to gather information about which states might have been visited. To illustrate this concept, we examine an example where sequences evolve in codon space and are observed in amino acid space. For simplicity, we disregard transition-transversion bias (i.e., we assume *κ* = 1). Given an alignment and phylogeny that strongly suggest an evolutionary trajectory W → L → H, the most direct path through codon space in single nucleotide steps requires at least two synonymous substitutions (Figure 1). Hence, knowledge of amino acid sequence evolution can reveal information about codon changes. In practice, many different routes through codon space may be compatible with the observed data; each is assessed by standard likelihood calculations and the embedded information about codon changes weighted appropriately.

**Fig. 1.**
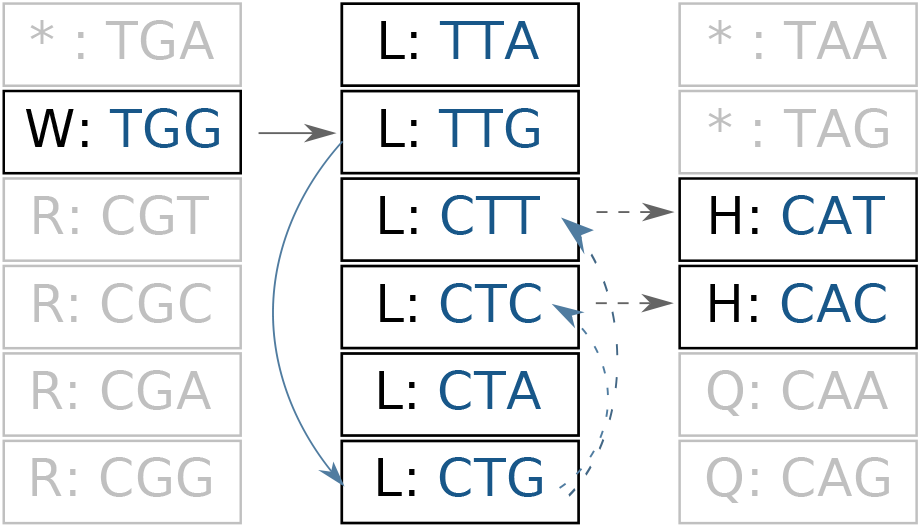
The path of a sequence through amino acid space contains information about which codons may have been visited. We illustrate a W (tryptophan) *→* L (leucine) *→* H (histidine) trajectory that requires multiple synonymous substitutions. Amino acid states are shown in black. Compatible corresponding (unobserved) codons are shown in blue. Solid arrows indicate changes required for the trajec tory with the minimum number of steps, while dashed arrows indicate substitutions associated with multiple compatible paths. Black arrows denote non-synonymous substitutions, and blue arrows indicate synonymous substitutions.

Where the separate state-space model disallows many transitions (as with the codon model in equation 1), it is easy to see how inferred moves through the aggregated space can give information about the separate states. However, even when the model of interest in **S** is described by a *Q*-matrix that does not contain zeros, similar principles apply. Here, none of the routes through the unobserved space are prohibited by the exchangeabilities, but each is more or less probable. We can therefore distinguish between different routes without directly observing them. For example, given an alignment that implies the amino acid trajectory L *→* F *→* Y, the RAM55 model has several available routes through rotamer space. However, considering the relative empirical exchangeabilities between states, we observe marked differences in how probable each path is (Figure 2). This, in turn, should allow us to infer, for example, the most probable rotamer sequence at an ancestral node, using the ambiguity approach.

**Fig. 2.**
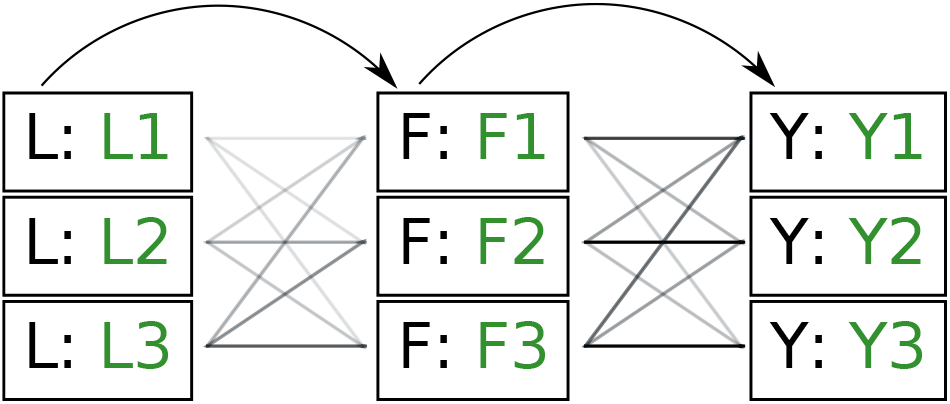
Illustration of paths through amino acid and rotamer space, given an implied trajectory L (leucine) *→* F (phenylalanine) *→* Y (tryptophan). Black font indicates observed amino acid states. Green font indicates unobserved rotamer configurations. Arrows show observed path through amino acid space. Lines connecting rotamer states indicate transition probabilities between states, with darker shading indicating more probable substitutions according to the RAM55 matrix. L3 *→* F3 *→* Y3 is the most likely trajectory.

### Selection can be inferred from amino acid data alone

Next, the question arises whether ambiguity coding extracts enough signal to allow meaningful inferences to be made. We therefore asked whether the ambiguity approach permits inference about codon evolution from amino acid data, considering the M5 variant of the standard M0 codon model (see Methods). To determine if M5 is capable of detecting the relative strength of selection under which a sequence evolved, we require data for which this parameter is known. The most straightforward way of obtaining this is to simulate sequences under a model identical to the one used in the estimation step.

We therefore considered translated sequences that were evolved on a randomly generated 20-taxon tree under the codon model M0. As an initial benchmark, we generated 100 alignments with 3000 codons with the simulation parameters *ω*\* = 0.3 and *κ*\* = 2, and obtain accurate and unbiased estimates of both (median 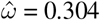; median 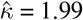). Analysing the original codon sequences using M0 with identical settings gives similar results (median 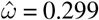; median 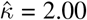). We note that M5 tends to be noisier, presumably due to its inability to directly observe synonymous changes. This observation holds across a range of *ω*\* and *κ*\* values, with high *ω\** values being somewhat prone to overestimation, although a strong linear relationship between true and estimated parameters is maintained (Figure 3). There is little interaction between *ω* and *κ* (Figure S3). In the following, we therefore consider only the combination *ω*\* = 0.03 and *κ*\* = 2, unless otherwise noted.

**Fig. 3.**
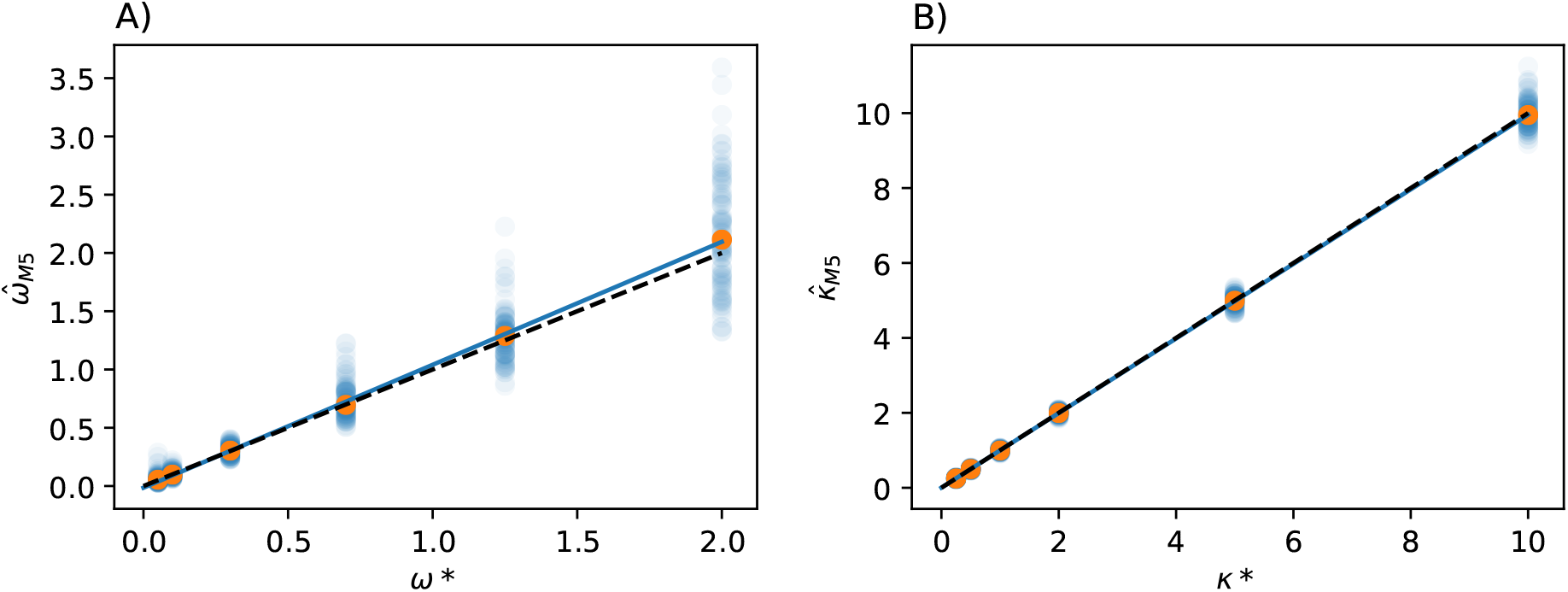
Estimation of codon model parameters from amino acid data. **A)** Simulation (*ω*^∗^) and estimated 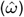 values of *ω* show a strong linear relationship (*κ*^∗^ = 2 for all simulations shown). Orange points represent the median 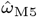 for each *ω*^∗^; blue points show estimates from individual alignments. Dashed line indicates *y* = *x*; solid blue line shows the line fit for the medians, with high values of *ω*^∗^ slightly prone to overestimation (slope = 1.07, intercept = 0.0083, *r*^2^ = 0.92, *p* ≈ 0). **B)** Estimates of *κ*^∗^ show a similar pattern (*ω*^∗^ = 0.3; slope = 1.00, intercept = −0.0009, *r*^2^ = 1.00, *p* ≈ 0).

The parameter 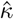 shows a relatively modest increase in variance under M5 compared to M0 (standard deviation *s*_*κ*_ = .0541 under M5; *s*_*κ*_ = 0.0364 under M0), presumably due to its direct, and thus inferable, impact on non-synonymous substitution patterns. The results obtained for 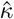 under M5 are similar to those for M6 (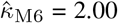 with *s*_*κ*_ = 0.0561), which estimates the parameter from amino acid sequences by averaging over synonymous codons rather than gathering information from ambiguity coding while traversing the tree [1]. This suggests that 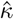 can be robustly estimated from amino acid sequences, even using coarse-grained approaches.

However, as expected, discarding codon information does lead to a loss of signal primarily affecting 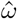, which displays a markedly higher variance under M5 than under M0 (*s*_*κ*_ = 0.0456 *vs.* 0.0056, respectively). Why might this be? A comparison of the estimates for *dS* tree length versus *dN* tree length suggests that M5 has more difficulty estimating the former, with variation in *dS* tree length accounting for almost all of the variation in overall tree length (see Figure S4). This is consistent with the fact that synonymous changes are not directly represented in amino acid sequences, whereas non-synonymous changes are (as long as sufficiently short timescales are considered). We therefore next consider how much information about *ω* is retained by M5, compared to M0.

### How much information loss does discarding codons cause?

The ability of M5 to capture information that is directly ‘seen’ by M0 can be measured by comparing the variances in parameter estimates on alignments with varying amounts of evolutionary signal. The most straightforward way to add information to a phylogeny given a codon model is to increase the number of codons in the alignment. Since both M0 and M5 give rise to unbiased estimates of 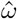 and 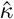 (Figure 3), we compared the variance in the parameter estimates for alignments of varying lengths. Given *ω\** = 0.3 and *κ\** = 2 across 1000 replicates, we find that the median standard error of 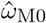, as estimated by codeml, is consistently lower than that of 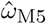. Across the range, the standard error is approximately 10 times higher for M5 (Figure 4), indicating an information loss of 98.7% (see Figure 4A for details). By comparison, the equivalent loss for 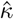 is only 38.8% (Figure 4B). Nevertheless, the estimates of 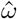 are reasonably accurate given a sufficiently long alignment (Figure 3).

**Fig. 4.**
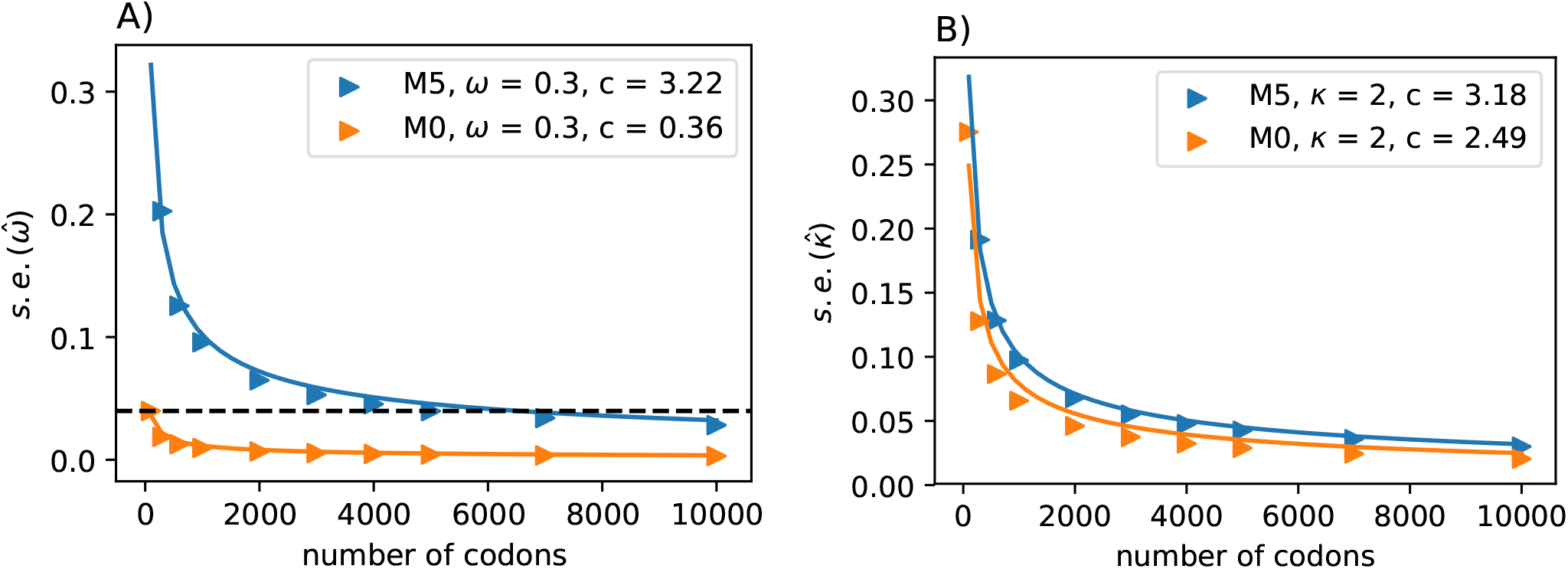
Increasing the number of columns allows us to quantify how much information is retained in amino acid sequences (M5) relative to codon sequences (M0). The median standard errors of **A)** 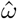 and **B)** 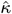 decrease for both models as codons are added. The dashed horizontal line in A) indicates the observed median standard error of 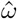 for 100 codons under M0, and illustrates that M5 requires a substantially longer alignment to reach a comparable standard error. Fitting functions of the form 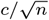 to the median standard errors, with *n* equalling the number of alignment columns, allows us to quantify the difference in information content. Equating 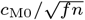 with 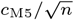 indicates equivalent information content for *fn* codons in M0 and *n* codons in M5: hence M5 recovers a fraction *f* = (*c*_M0_/*c*_M5_)^2^ of the information available to M0. Alternatively, M5 has lost 100(1 *−f*)% of the information available to M0.

It is perhaps counter-intuitive that acceptable estimates of 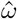 can still be obtained with M5 in these circumstances. However, the relative magnitude of the error may remain relatively small. For example, given *ω* = 0.3, *κ* = 2 and *n* = 3000, the interquartile range for 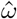 is 0.296 - 0.304 under M0 and 0.270 - 0.344 under M5. Note that the information loss seems to vary with *ω*^∗^. We observe a larger ratio of standard errors when *ω*^∗^ = 0.1 (approximately 12–20 times), with an information loss of about 99.2% (results not shown).

### Tree depth and taxon number impact M5 information loss

Our results show estimates of 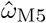 to be noise-prone for short alignments. Because increasing alignment length is not a viable solution to reduce variance for real amino acid sequence data, we also considered how other features of the alignment impact M5’s ability to accurately infer 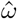. For example, it may be possible to select sequences with higher divergence or include additional taxa in the phylogeny, hence adding more information. Scaling-up the branch lengths of the tree in the simulations gives an initial improvement in the standard error of 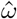. The greatest reduction is observed for a tree length of approximately 4 times the length of the original tree (around 20), followed by a decline for longer trees (see Figure 5). This is a consequence of the increased number of substitution events from which the model can infer parameters, which is advantageous until the sequences become too divergent, the true number of substitutions is underestimated, and the data become too noisy.

**Fig. 5.**
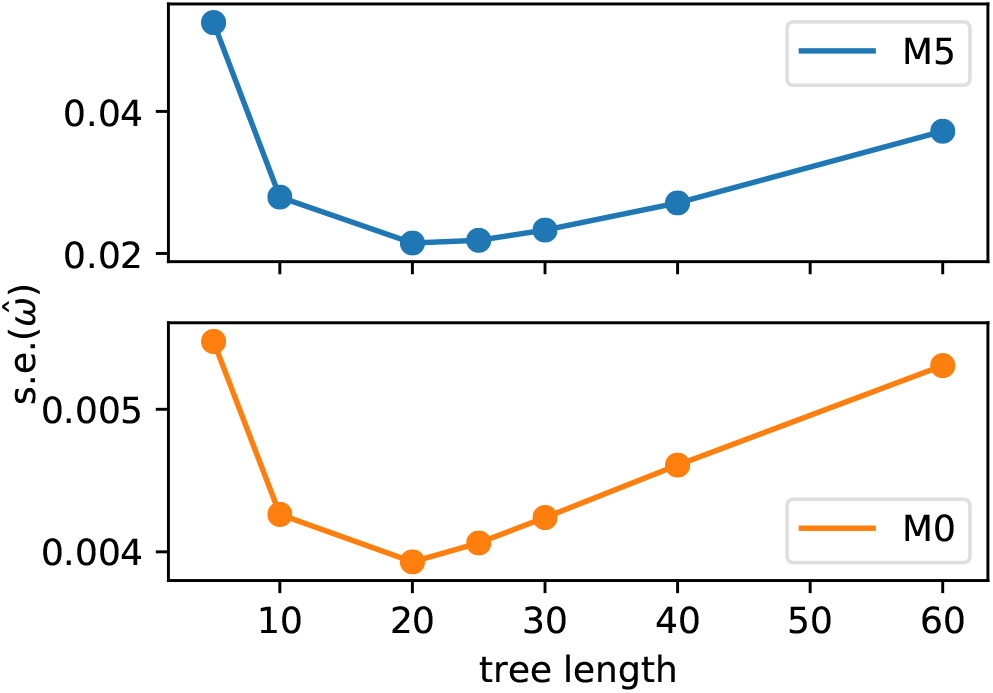
Tree length influences variance in estimates 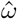 for both M5 and M0, with intermediate values producing the lowest standard errors. Points show the median value across replicates. Note the difference in scale on the y-axis between M5 and M0.

The trajectory of the change in variance observed across different tree lengths is broadly comparable for M0 and M5, with the variance for M5 remaining consistently higher for all lengths. In the case of M0, this is due to saturation at synonymous sites [2]. For M5, it is easy to see that multiple substitutions at a site along a single branch make it more difficult to infer the path through codon space (see Figure 1). This confirms that the model is behaving as expected.

Adding additional taxa has a similar effect on the variance. When we examine a tree of comparable height (0.5) with twice as many tips (*n* = 40, see Figure S1B), the standard error of 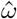 decreases compared to the smaller tree (median = 0.0223 vs .0.0396 for 1000 replicates of 5000 codons and *ω*^∗^ = 0.3; equivalent values for M0 are 0.0030 vs. 0.0043). As above, this behaviour is expected as the additional tips add information, provided the branches are not exceedingly long.

Given these observations, we conclude that estimating the strength of selection from amino acids is a feasible strategy where nucleotide sequences may be difficult or impossible to obtain. Although there is an appreciable loss of signal, M5 is statistically consistent and approaches the correct parameter estimates given enough amino acid sequence data.

### Accurate reconstruction of ancestral side chain configuration from amino acids

We next ask whether the strategy of treating characters that are not directly observable as ambiguous is also informative when the instantaneous rate matrix underlying the substitution model is not sparse (that is, does not contain transitions with probabilities equal to 0). To examine how ambiguity coding performs given an empirical rotamer-aware model, we simulated data with 55 states under the RAM55 model on a 32-taxon tree (Figure S2, see Methods), and subsequently reconstructed the ancestral sequences under the same model. We opted to benchmark the model using reconstruction accuracy, as ancestral side chain configurations represent an output that would be otherwise unobtainable from amino acid data alone. Varying the proportion of masked sequences in the alignment (see Methods) allows us to compare scenarios where structures are available for some of the sequences of interest, or none at all, similar to what would be observed for real empirical data. The reconstruction accuracy for the data where rotamer information is available for all of the tips provides a benchmark for the performance of ambiguity coding.

There is a relatively modest reduction in overall rotamer state reconstruction accuracy between simulations where rotamer configurations are known for all taxa, and simulations where this information is not available for any of the taxa (~ 15% difference for the unscaled tree, Figure 6, scaling factor = 1.0). Reconstruction under RAM55 using only amino acid sequences is markedly more accurate than the only alternative approaches of using a conventional empirical amino acid model to reconstruct the protein sequence and randomly assigning (‘guessing’) rotamer states (Figure 6, dashed red line), or assigning them based on the equilibrium frequencies of the RAM55 model (Figure S5). Hence, it is advantageous to reconstruct under the rotamer-aware model, even when the input data are only available in the aggregated state-space.

**Fig. 6.**
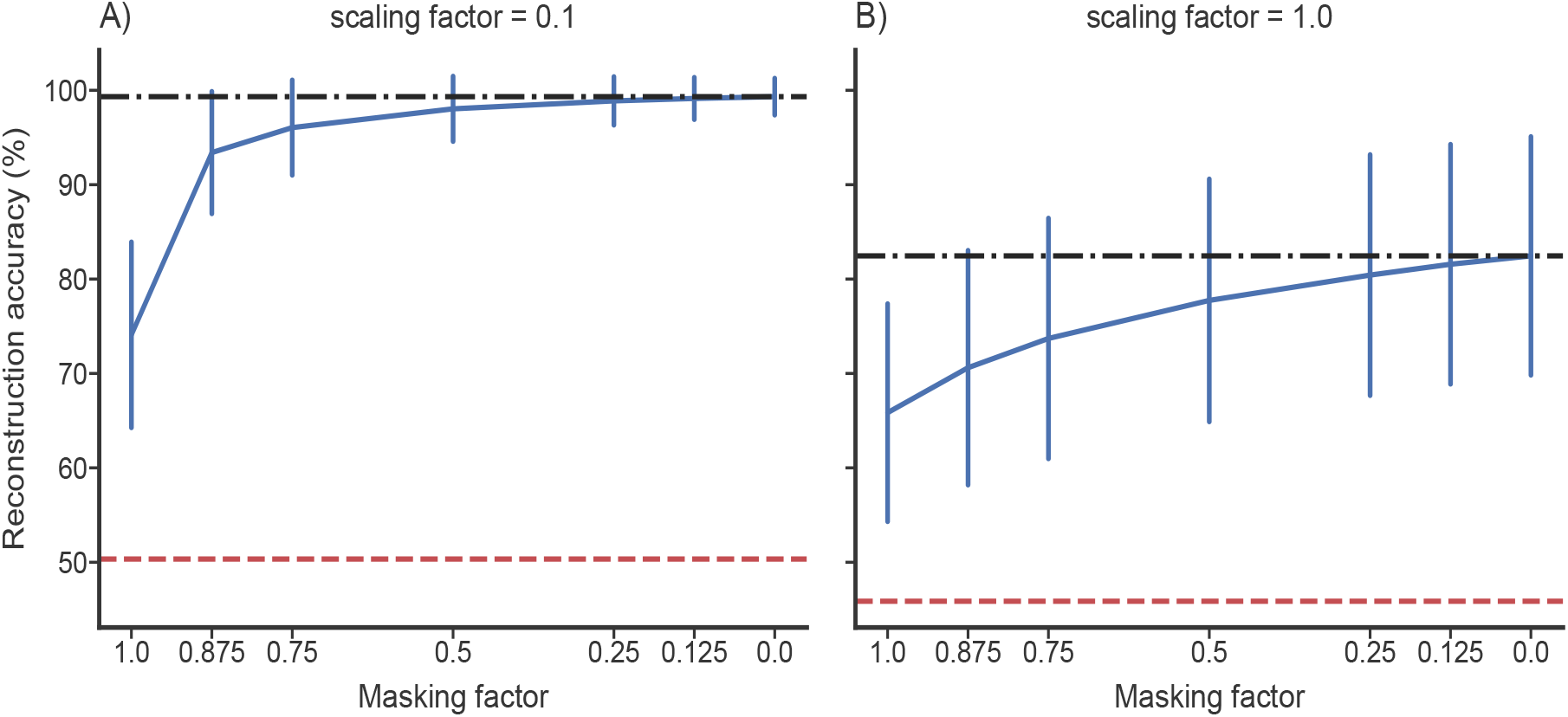
The accuracy of ancestral rotamer sequence reconstruction from mixed data under RAM55 increases and shows lower variance when more rotamer configuration information is available. The x-axis shows the fraction of rotamer configuration information removed (i.e. masked). The vertical bars show the standard deviation of the reconstruction accuracies, centered around the median. The black dash-dot lines represent the maximum accuracy reached on full (unmasked) alignments; red dashed lines show the accuracy achieved by reconstructing the amino acids under RUM20 and randomly assigning (‘guessing’) the rotamer configuration. **A)** Results when all branches of the tree (Figure S2) are multiplied by 0.1, showing greater overall accuracy in this case. **B)** Results for the standard simulation tree (see Methods).

As expected, the overall accuracy depends on how difficult the ancestral sequence reconstruction problem is, with a shallower tree showing higher sequence identity between simulated and reconstructed characters (Figure 6A). Interestingly, the greatest increase in performance appears between alignments with no rotamer configuration information present and 12.5% of sequences containing that information (Figure 6). This suggests that little structural information is required in order to achieve ancestral reconstruction of rotamer states with acceptable accuracy. Intuitively, the fraction of correctly inferred states declines with increasing distance from the tips of the tree (Spearman’s rank correlation coefficient −0.537, *p* < 0.001; details not shown). We also considered how the certainty with which the model assigns the correct ancestral state responds to rotamer information being masked at the tips of the tree. Unsurprisingly, the marginal posterior probability for the correct state declines as information is removed (see Figure S6). We observe a drop in the certainty of the reconstruction preceding the drop in accuracy.

### Gains associated with using amino acid sequences to infer rotamer configuration in absence of structure

As with the codon model example, we would like to quantify the loss of information associated with using aggregated state-space data for inference in the separate state-space. Given that the output of the empirical model we are studying is not a parameter estimate (as opposed to our mechanistic codon model/selection example) but the percentage of correctly reconstructed residues, extending the alignment is not informative. Instead, we compared the accuracy of reconstructions under two scenarios: (a) all state information (amino acid and rotamer configuration) is discarded from a proportion of sequences, and (b) masking is used so that amino acid, but not rotamer, sequences are available for a proportion of the alignment. This provides a measure of the advantage gained by considering additional amino acid sequences where no structural information is available.

For the un-scaled tree, masking 50% of the rotamer configurations produces ancestral reconstructions that are comparable in accuracy to trees where 12.5% of taxa have gaps (Figure 7A), indicating a noticeable advantage for including amino acid sequences where full rotamer state information is unavailable. In other words, augmenting half of the amino acid sequences with rotamer configuration information is approximately as informative as having 87.5% of the full rotamer information. Further, removing all rotamer information and reconstructing with ambiguity is equivalent to retaining 50% of the original information. These results suggest that it can be very valuable to consider amino acid sequences that lack structures.

**Fig. 7.**
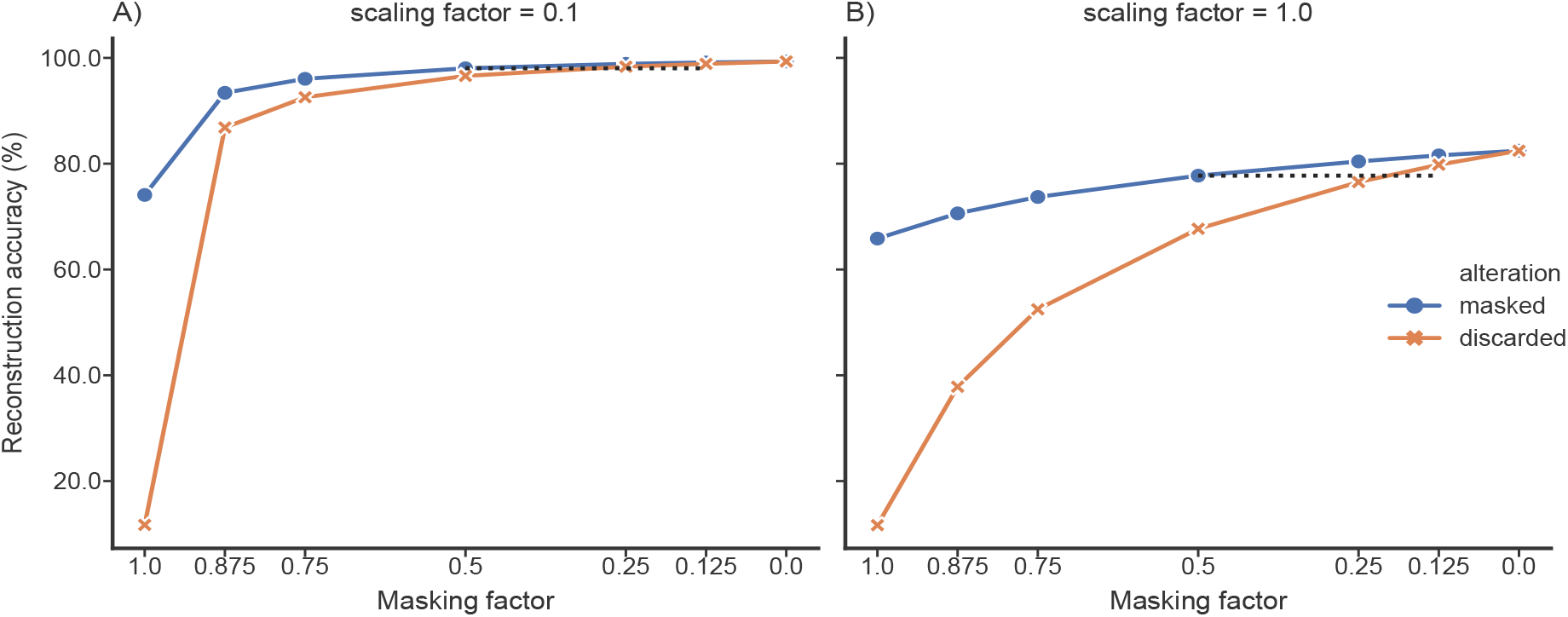
The accuracy of ancestral rotamer sequence reconstruction from mixed data under RAM55 increases when masked sequences, which lack rotamer states, are not discarded. The x-axis shows the fraction of information available under two scenarios. The blue points reflect the the amount of rotamer information that has been masked (i.e. replaced with amino acids), and the orange points represent rotamer sequence that has been replaced with gaps (discarded). Masking half of the rotamer configurations produces accuracies comparable with those obtained by replacing 1/8–1/4 of sequences in the alignments with gaps (see black dashed lines). Reconstructions with ambiguity always outperform discarding an equivalent fraction of amino acid sequences. As before, the shallower tree, A), shows higher overall accuracy.

### Improved prediction of side chain configurations in homologous structures

Considering its robust performance, how might ambiguity coding be put to practical use in the context of reconstructing side chain configurations? Prediction of side chain conformations is an important part of protein structure modelling and interaction modelling. For a given protein sequence of unknown structure, it is possible to construct a model of the target protein from its amino acid sequence and experimentally determined structures of related homologous proteins. This homology modelling strategy aims to predict both the main chain geometry and side chain configurations. In conserved regions, side chains can be modelled starting from configurations observed at corresponding sites in the nearest homologous structure (NNC: see Methods). Further steps are then required, particularly to model non-conserved side chain configurations [10]. Side chain configurations could be predicted for an extant amino acid sequence using RAM55 and a modified ancestral reconstruction algorithm by constraining the *χ*_1_ configuration prediction to the set of configurations that are possible given the observed amino acid at any given site. Another realistic homology modelling scenario might involve our target’s nearest homolog also lacking a resolved structure and only being available as an amino acid sequence (MNN: see Methods). In this context RAM55 can use a mixed alignment (amino acid sequences and rotamer sequences) to inform its predictions rather than relying exclusively on available structures, which ought to improve reconstruction accuracy as seen above (Figure 7).

To evaluate our approach to side chain configuration prediction, we considered two empirical protein family datasets (RuBisCO and ADK, see Methods) composed of amino acid sequences from a range of species and their corresponding rotamer configuration information. We investigated two scenarios: (1) a rotamer sequence is available for the nearest neighbour of each terminal node or (2) only a masked amino acid sequence is available for the nearest neighbour. Predicting *χ*_1_ side chain configurations using RAM55 is more accurate (~ 11% median improvement) than NNC (see Methods) when the nearest neighbour’s structure is available (Figure 8; see also Supplementary Figures S7, S8). Further, RAM55 can make use of all the available rotamer sequence information, as well as the nearest neighbour’s amino acid information, when the nearest neighbour’s structure is not available. Meanwhile, the traditional approach would instead rely on the second-nearest structure (MNN: see Methods). This results in improved reconstruction accuracy for RAM55 (~9% median improvement) over MNN (Figures 8, S7, S8). For both NNC and MNN analyses, the improvements with RAM55 are driven by strongly increased accuracy at nonconserved sites (results not shown).

**Fig. 8.**
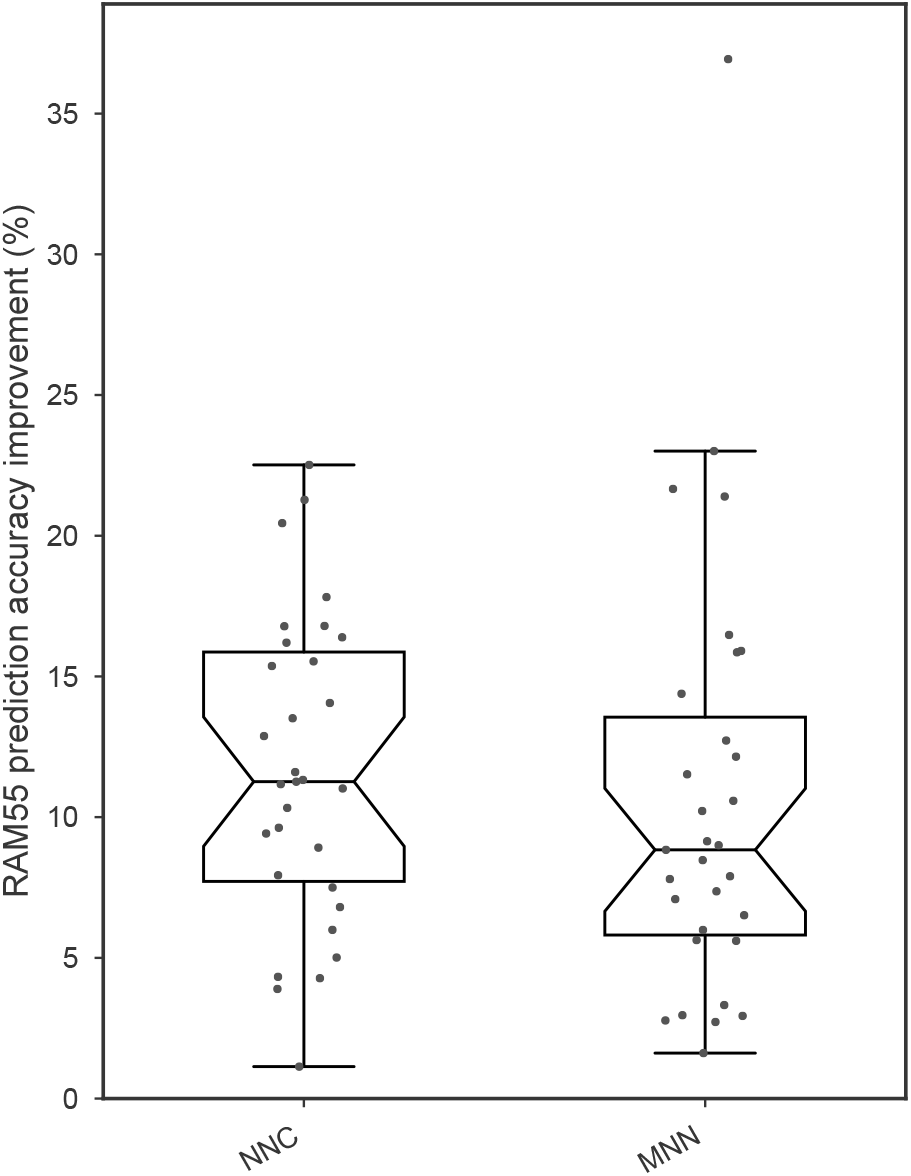
Improvement in side chain configuration prediction from different strategies that incorporate structural information and sequence. RAM55 predictions make use of all the information available in the RuBisCO dataset (mixed amino acid and rotamer sequences) where the nearest neighbour of each predicted sequence is available as a rotamer sequence (for NNC) or is masked (for MNN). Accuracy improvement is computed against scores from two different strategies, based on the nearest neighbour’s configuration or the second-nearest configuration (NNC or MNN, respectively; see Methods). RAM55 achieves a higher accuracy in both cases (Wilcoxon one-sided signed-rank tests both *p* < 0.001).

Our method provides plausible predictions of *χ*_1_ configurations using a strategy that, as opposed to NNC or MNN, explicitly models the evolutionary process along the branches of the phylogeny and can make use of amino acid information when structures are not available. RAM55-based predictions could speed up the side chain homology modelling process by creating an informed prior to constrain the search space, particularly where close homologs with unresolved structures might otherwise be discarded by traditional strategies.

## Discussion

We have demonstrated that treating characters in an aggregated state-space **A** as ambiguous versions of characters in a larger state-space **S** allows us to obtain information that would otherwise not be accessible from data in **A**. Our examples show that this is true for estimating the strength of natural selection under a codon model, and for reconstructing ancestral side chain configurations under an empirical model, both from amino acid sequences alone.

Naturally, where data are available in a larger state-space matching the internal structure of the preferred model it is advantageous to make use of them. The codon model example provides a particularly clear illustration: completely discarding codon information leads, for obvious reasons, to increased variance in estimates of 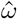. We nevertheless find it remarkable that selection parameter estimates ordinarily derived from comparisons of synonymous and nonsynonymous substitutions can be obtained given sufficient amino acid data.

It has previously been argued that modelling coding sequence evolution at the codon level rather than the amino acid level is generally preferable because it offers a more detailed description of the process that generated the data [3, 27–30]. On the other hand, the ambiguity approach may be useful to obtain an approximate estimate of the strength of natural selection in cases where amino acid sequences are more readily obtainable. For example, the supplementary materials accompanying phylogenetic studies often only provide amino acid alignments, and as many as 17% of nucleotide sequences corresponding to proteins in Pfam [31] have been previously reported unrecoverable [32]. The ability to perform a preliminary screen to determine whether a sequence of interest is under weak or strong evolutionary constraint might therefore be convenient. However, we caution against over-interpreting the results returned by M5, particularly when individual sequences are being considered or codon usage may be biased.

The absence of high-quality structures for many extant proteins provides more practical applications for ambiguity coding. In the case of the RAM55 empirical rotamer model, we have shown the utility of using amino acid sequences alone, and ‘mixed’ inputs where even a limited amount of structural information leads to considerable improvements in the accuracy of *χ*_1_ configuration prediction. Being able to use information from amino acid sequences improves prediction accuracy over modelling side chains based on the nearest available structure alone. This approach could benefit homology modelling strategies, specifically the steps involving modelling both conserved side chains based on a known template structure, and non-conserved side chain modelling achieved by searching a rotamer library and minimizing an energy function [33–35]. In this context, RAM55’s predictions constrain the rotamer configuration sampling space. This could result in a reduction of the number of energy refinement cycles required.

In addition, using RAM55 and the marginal ancestral reconstruction algorithm makes it possible to obtain posterior probabilities for each of the possible configurations at a given site. This distribution might provide a more-robust prior for further refinement, compared to using the single most-likely reconstructed configuration or the nearest homolog’s configuration at that site. Further work would be required to quantify improvements in speed and accuracy.

Given the advantage of including mixed input data demonstrated in our rotamer sequence reconstruction analyses, we expect combining amino acid and DNA sequences to be promising, as well as straightforward to implement. This would address some of the current limitations of M5 with respect to analysing phylogenetic datasets with some missing codon sequences. A more speculative and potentially intriguing application would be estimating selection or structural information from ancient protein sequences. Proteins can persist for longer in the environment than DNA under certain conditions [36–38], enabling phylogenetic inferences to be made based on substantially older specimens such as dinosaurs [39,40]. Our methods permit the use of a mixture of all available DNA and protein sequences to maximise signal, extending analyses that are normally only possible with DNA sequences to incorporate additional data sources. In the absence of any compelling available ancient protein datasets, we do not attempt to provide a benchmark here.

In this work, we have shown that ambiguity coding allows evolutionary inference from partially ‘hidden’ data under phylogenetic models with both sparse (e.g. mechanistic) and non-sparse (e.g. empirical) exchangeability matrices. Thus, the principles underlying likelihood analysis of missing data [2,12] and covariotide models [11] can be applied more broadly, allowing us to estimate selection and reconstruct aspects of protein structure given input data that are not fully resolved. Finally, ambiguity coding could conceivably be applied to other state-spaces beyond amino acids, codons, and rotamer states, provided there is reason to believe that movement through the aggregated space contains info about the separate space.

## ACKNOWLEDGEMENTS

We thank Nicola de Maio, Iain Moal and Alexey Kozlov for helpful discussions.

## MODEL AVAILABILITY

The M5 codon model is available as model 5 in codeml (PAML version 4.9h) [16] and is run with the sequence type set to amino acids (seqtype = 2). The program overrides the codon frequency setting specified in the control file and resets the CodonFreq variable to 0 (1/61). Rotamer sequence simulation and ancestral sequence reconstruction code is available at https://bitbucket.org/uperron/ambiguity_coding.

