## Supplementary Material for "Ambiguity coding allows accurate inference of evolutionary parameters from alignments in an aggregated state-space"

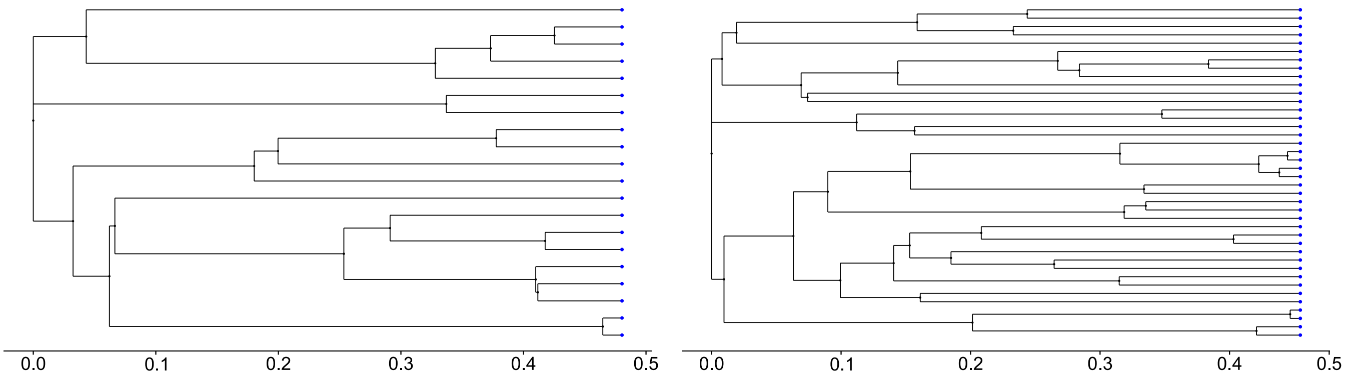


**Supplementary Figure 1. A)** 20-taxon birth-death tree used for codon simulations. (**B)** 40-taxon birth-death tree used for codon simulations. Tree height is shown on the x-axis.

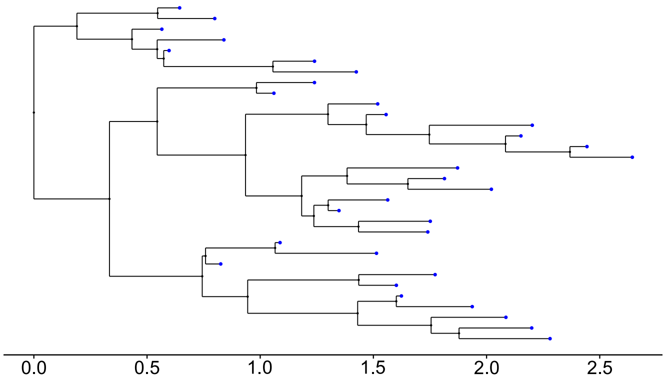


**Supplementary Figure 2.** 32-taxon Yule process tree used for rotamer simulations. Tree height is shown on the x-axis.

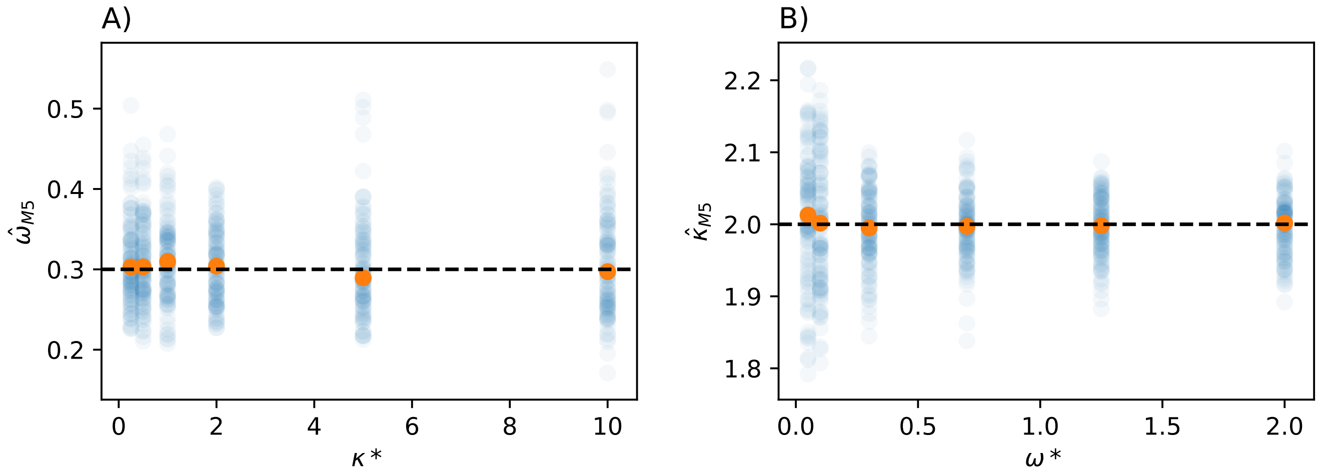


**Supplementary Figure 3. A)** M5 $\hat{\omega}$ estimates from data simulated with $\omega^*$ = 0.3 and different values of $\kappa^*$. **B)** M5 $\hat{\kappa}$ estimates for data simulated with $\kappa^*$ = 2 and different values of $\omega^*$. $\hat{\kappa}$ is slightly overestimated and shows higher variance when $\omega^*$ is very small, relative to comparisons with an equivalent sequence length (3000 codons).

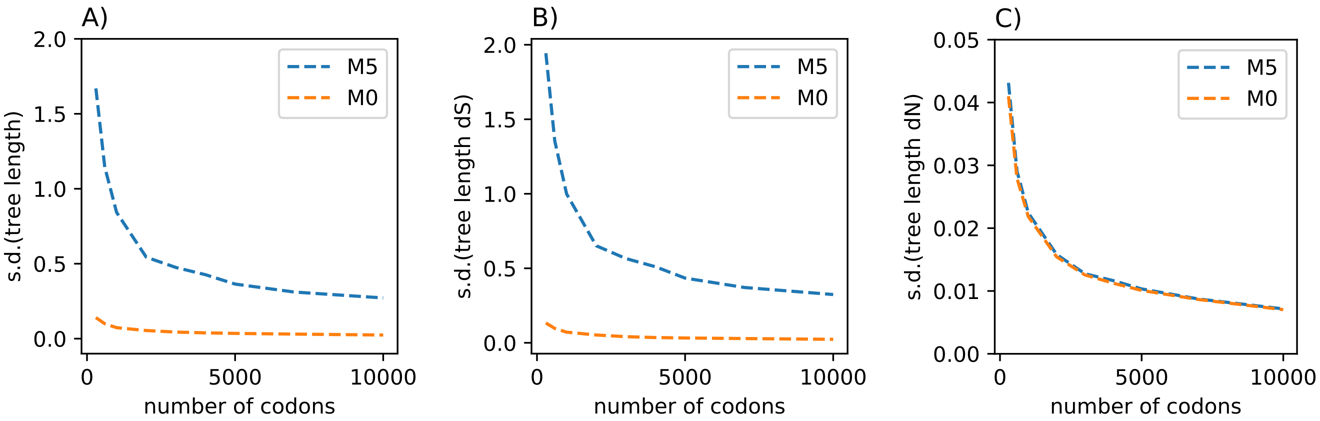


**Supplementary Figure 4.** The variance in the tree length estimates for M5 is almost entirely accounted for by *dS* tree lengths. Overall tree lengths **A)** and *dS* tree lengths **B)** for M0 are markedly less noisy than for M5. *dN* tree lengths **C)** show almost identical standard deviations across different alignment lengths. Data shown for $\omega^*$ = 0.3, $\hat{\kappa} = 2$.

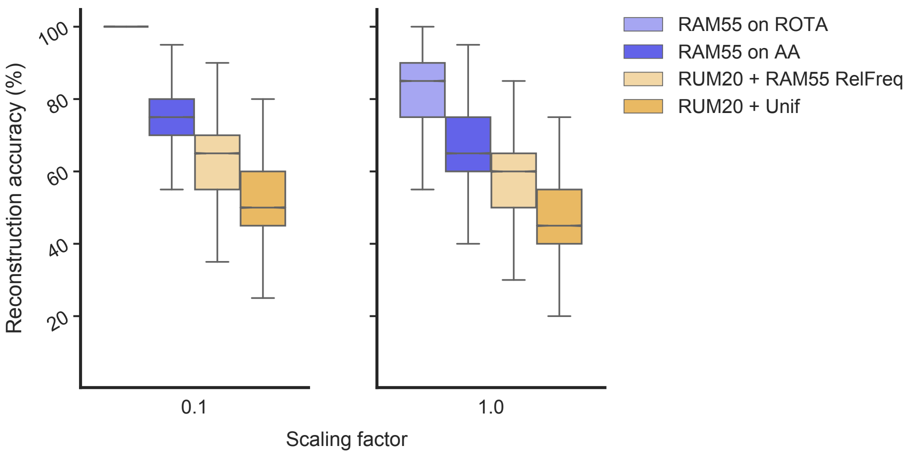


**Supplementary Figure 5.** Comparison of ancestral rotamer state reconstruction under the RAM55 55-state model and random assignment of states. Ancestral states are reconstructed from an alignment of 32 simulated sequences from a fixed tree (see Figure S2) whose branches are scaled according to a factor reported on the y axis. ‘RUM + Unif’ refers to amino acid sequence reconstructions that were assigned rotamer states at random (based on the assumption of equal frequencies), while ‘RUM20 + RAM55 RelFreq’ were assigned rotamer states based on the equilibrium frequencies of RAM55. These assignments all fare worse than reconstruction under RAM55 from either amino acid (‘RAM55 on AA’) or (‘RAM55 on ROTA’), indicating that the information retained is attributable to our model. For the first panel the (‘RAM55 on ROTA’) reconstructions are correct across all sites and replicates.

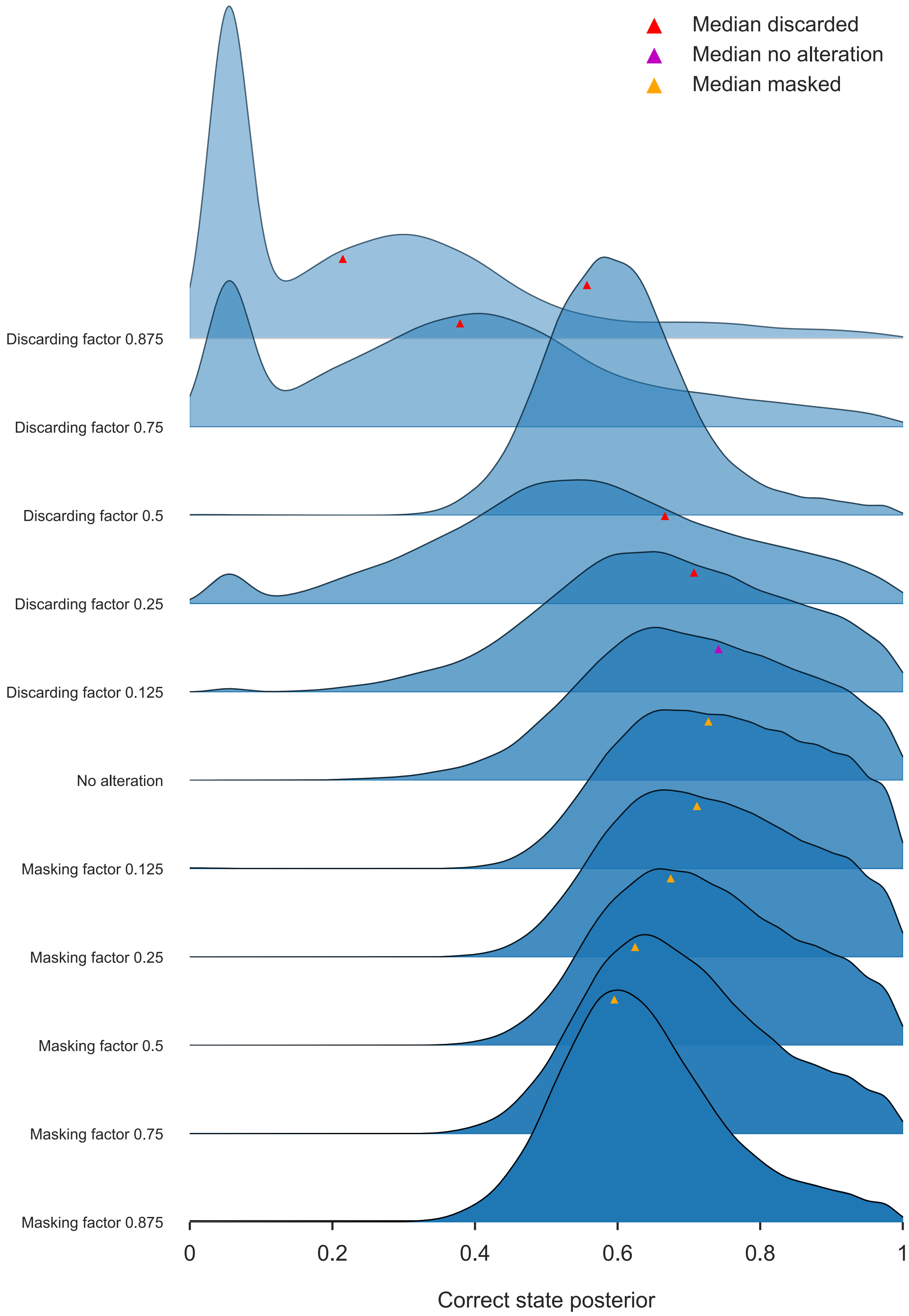


**Supplementary Figure 6.** Model certainty, as measured by the marginal reconstruction's posterior, improves when more rotamer configuration or amino acid information is available. For any proportion of the 32 sequences (Figure S2, scaling factor = 1), including masked amino acid sequences (orange), rather than discarding them (red), increases the posterior probability assigned by the model to the correct ancestral state. As more information is removed, decrease in certainty precedes the decrease in accuracy (see Figure 7). Each subplot reports the kernel density estimation for the posterior probability of the correct ancestral state, medians for each posterior distribution are highlighted.

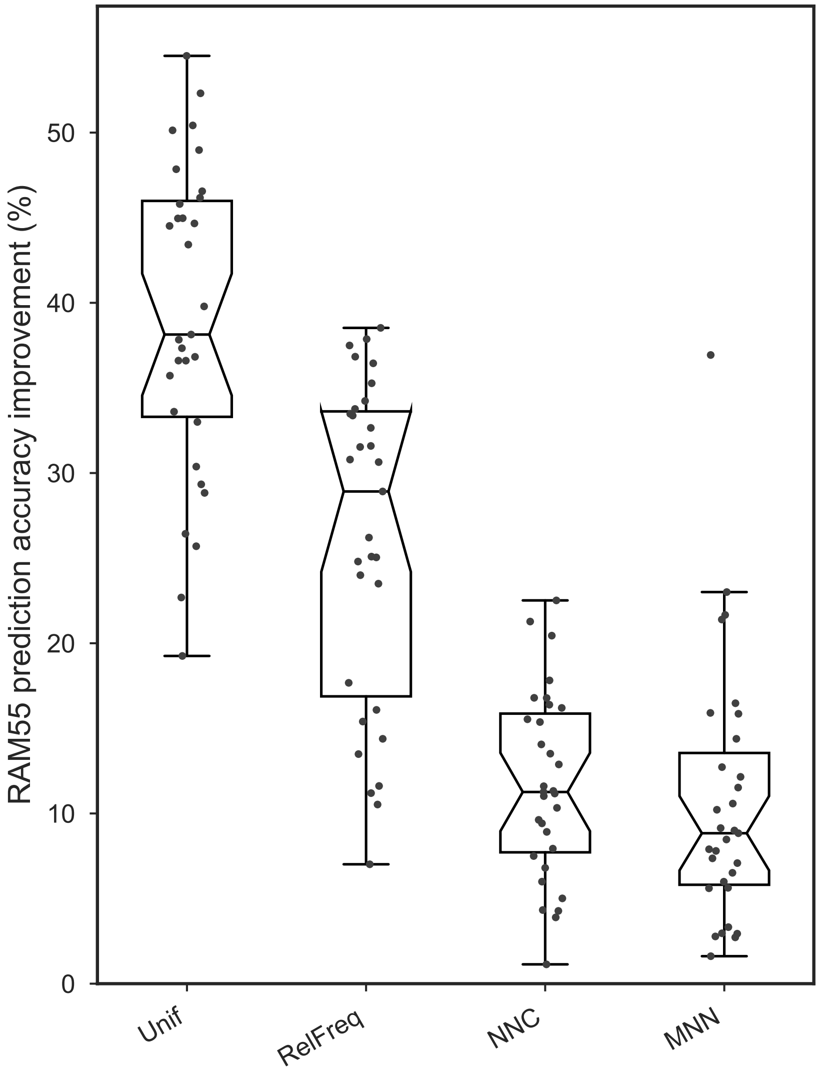


**Supplementary Figure 7.** Improvement in side chain configuration prediction from different strategies that incorporate structural information and sequence on the RuBisCO empirical dataset (31 taxa). We iterate over all terminal nodes and reconstruct each amino acid's $\chi_1$ configuration using RAM55 and the marginal algorithm while constraining the reconstruction to the states possible given the observed amino acid. Accuracy improvement is computed against four configuration assignment strategies: a uniform distribution (Unif), the relative RAM55 equilibrium frequencies of each configuration for the observed amino acid (RelFreq), assignment based on the nearest neighbour's configuration (NNC), and MNN, which is like NNC but uses the second-nearest structure. Percentage of improvement in reconstruction accuracy is shown on the y-axis.

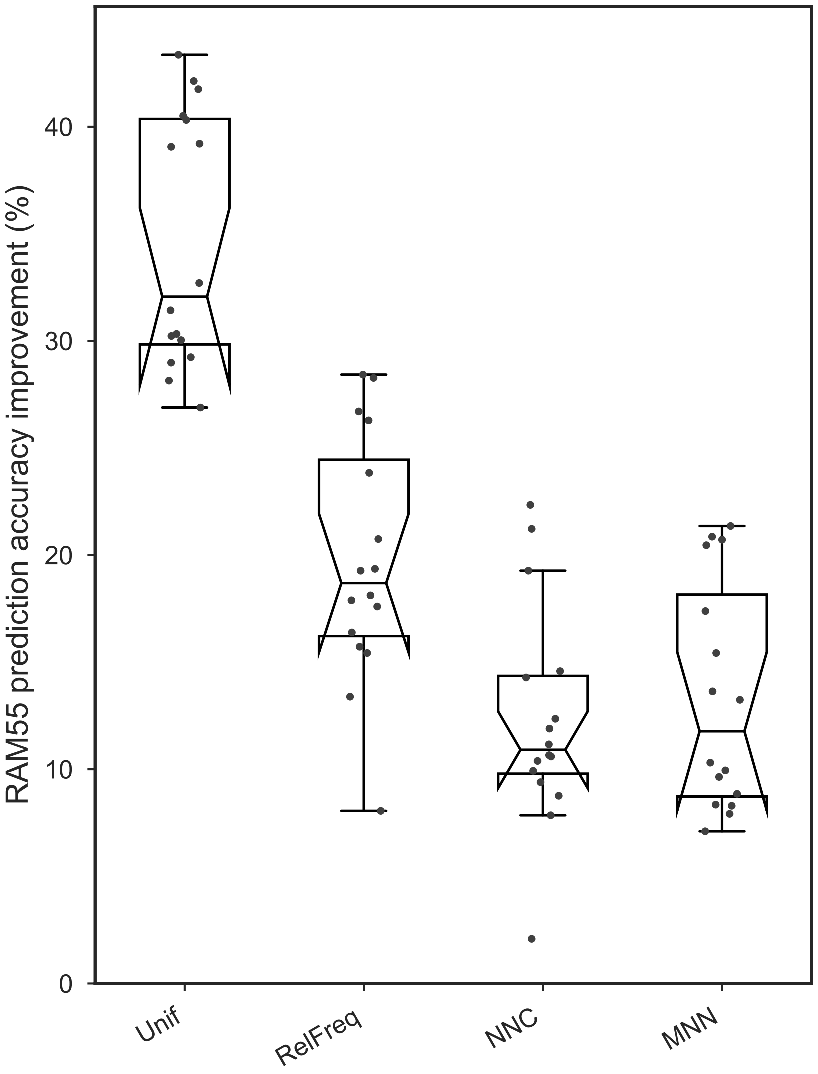


**Supplementary Figure 8.** Improvement in side chain configuration prediction from different strategies that incorporate structural information and sequence on the ADK empirical dataset (16 taxa). We iterate over all terminal nodes and reconstruct each amino acid's $\chi_1$ configuration using RAM55 and the marginal algorithm while constraining the reconstruction to the states possible given the observed amino acid. Accuracy improvement is computed against four configuration assignment strategies: a uniform distribution (Unif), the relative RAM55 equilibrium frequencies of each configuration for the observed amino acid (RelFreq), assignment based on the nearest neighbour's configuration (NNC), and MNN, which is like NNC but uses the second-nearest structure. Percentage of improvement in reconstruction accuracy is shown on the y-axis.
